# Synchronized β cell response to glucose is lost concomitant with loss of islet architecture in Robo deficient islets of Langerhans *in vivo*

**DOI:** 10.1101/2019.12.11.873471

**Authors:** Melissa T. Adams, Christopher A. Reissaus, JaeAnn M. Dwulet, Erli Jin, Joseph M. Szulczewski, Melissa R. Lyman, Sophia M. Sdao, Sutichot D. Nimkulrat, Suzanne M. Ponik, Matthew J. Merrins, Richard K.P. Benninger, Raghavendra G. Mirmira, Amelia K. Linnemann, Barak Blum

**Affiliations:** Department of Cell and Regenerative Biology, University of Wisconsin-Madison, Madison, WI 53705, USA; Herman B Wells Center for Pediatric Research and Center for Diabetes and Metabolic Diseases, Indiana University School of Medicine, Indianapolis, IN 46202, USA; Department of Bioengineering and Barbara Davis Center for Diabetes, University of Colorado Denver, Anschutz Medical Campus, Aurora, CO. 80045, USA; Department of Medicine, Division of Endocrinology, Diabetes, and Metabolism, University of Wisconsin-Madison, Madison, Wisconsin 53705, USA; Kovler Diabetes Center and the Department of Medicine, University of Chicago, Chicago, IL 60637, USA

## Abstract

The spatial architecture of the islets of Langerhans is hypothesized to facilitate synchronized insulin secretion between β cells, yet testing this *in vivo* in the intact pancreas is challenging. *Robo βKO* mice, in which the genes *Robo1* and *Robo2* are deleted selectively in β cells, provide a unique model of altered islet spatial architecture without loss of β cell differentiation or islet damage from diabetes. Combining *Robo βKO* mice with intravital microscopy, we show here that *Robo βKO* islets lose synchronized intra-islet Ca^2+^ oscillations between β cells *in vivo*. We provide evidence that this loss is not due to a β cell-intrinsic function of Robo, loss of Connexin36 gap junctions, or changes in islet vascularization, suggesting that the islet architecture itself is required for synchronized Ca^2+^ oscillations. These results have implications for understanding structure-function relationships in the islets during progression to diabetes as well as engineering islets from stem cells.

## Introduction

The islets of Langerhans, which comprise the endocrine pancreas, are highly organized micro-organs responsible for maintaining blood glucose homeostasis. Islets are composed of five endocrine cell types (α, β, δ, PP, and ε) which, in rodents, are arranged such that the β cells cluster in the core of the islet, while other non-β endocrine cells populate the islet mantle, so that most β cells are in direct contact preferentially with β cells (homotypic interactions)^1^. Human islet architecture is more complex and, though its exact organization pattern is still debated, the prevailing idea is that it still follows a non-random distribution of the different endocrine cell types^2–6^. In agreement with this notion, computational analysis of human islet architecture found lower probability of heterotypic interactions and a higher probability of homotypic interactions between the various endocrine cell types than would be expected if the islet displayed random distribution of endocrine cells^7,8^. Thus, in both rodent and human islets, respective stereotypical islet architectures prioritize homotypic over heterotypic interactions between endocrine cell types^7^. The biological reason for preferential homotypic interactions between endocrine cells is not completely clear, but it has been suggested to be important for dictating the level of Connexin36 (Cx36)-mediated electrical β cell-β cell coupling, thus allowing for synchronization of glucose-stimulated insulin secretion (GSIS) between neighboring β cells^9,10^.

Activation of insulin secretion in the β cell is triggered when glucose from the blood enters the β cells through glucose transporters. As this glucose is metabolized, the ratio of intracellular ATP/ADP in the cells increases. This rise in ATP causes ATP sensitive K^+^ channels to close, resulting in membrane depolarization. The resultant depolarization causes voltage-gated Ca^2+^ channels to open, triggering an influx of Ca^2+^ into the cell, which in turn promotes exocytosis of insulin granules^11–13^. This chain of events is cyclical and thus results in oscillations of membrane potential, cytosolic Ca^2+^ levels, and insulin secretion in response to glucose^14^. Because β cells within an islet are gap-junctionally coupled, and thus electrically coupled, these oscillations are synchronous across all islet β cells^15^. It is thus hypothesized that preferential β cell homotypic contact allows for the necessary amount of gap junctions to form between neighboring β cells in order to synchronize the oscillations in an entire islet, facilitating pulsatile insulin secretion^10,16^. Indeed, modeling experiments in which the number of homotypic β cell-β cell nearest neighbor connections is lowered within an islet result in predicted perturbation of synchronous Ca^2+^ oscillations^10^. If this *in silico* prediction is correct, then disrupting spatial organization of the different endocrine cell types within the islet alone, without affecting any other property of the cells, would be sufficient to disturb synchronized insulin secretion between β cells. However, direct empirical evidence for this hypothesis is lacking.

Most genetic mouse models that show disruption of islet architecture also display defects in glucose homeostasis^17^. However, in many of these models, the disrupted islet architecture phenotype is linked to either developmental defects in β cell differentiation or maturation^18–29^ or to pathologies related to β cell damage in diabetes^30–35^. This introduces a strong confounding factor for studying the role of islet architecture on β cell function. Therefore, current mouse models of disrupted islet architecture are not suitable for directly testing the hypothesis that preferential homotypic β cell-β cell interactions, dictated by canonical islet architecture, regulate synchronized insulin secretion between β cells within the same islet.

Recently, we have described a mouse model in which the cell-surface receptors *Robo1* and *Robo2* are deleted specifically in β cells (*Robo βKO*), resulting in disruption of canonical endocrine cell type sorting within the islets^36^. Unlike other models of disrupted islet architecture, the β cells in the islets of *Robo βKO* express normal levels of markers for β cell differentiation and functional maturity. We reasoned that this model would allow for direct testing of the role of islet architecture on synchronous islet oscillations between β cells in a fully differentiated, non-diabetic islet setting.

## Results

### *Robo βKO* islets have fewer homotypic β cell-β cell contacts than control islets

*In silico* simulations where the degree of β cell-β cell coupling is changed through a decrease in homotypic nearest neighbors predict that disruption in islet architecture will disrupt synchronous intra-islet Ca^2+^ oscillations and hormone secretion pulses^7,10,37^. To test whether β cells in *Robo βKO* islets have a decreased ratio of homotypic β cell neighbors on average than control islets, we performed nearest neighbor analysis on islets from pancreatic sections of *Robo βKO* and control mice (**Figure 1**). We found that *Robo βKO* islets possess significantly fewer β cell-β cell contacts (*n*=9-11 islets for 3 mice from each genotype; control 75.35%±4.1, *Robo βKO* 50.37%±4.1, *p*=0.01), and homotypic contacts in general when compared to control islets (*n*=9-11 islets for 3 mice from each genotype; control 83.7%±1.7, *Robo βKO* 64.43%±1.2, *p*=0.0008). We also found that *Robo βKO* islets possess significantly more β cell-α cell contacts (*n*=9-11 islets for 3 mice from each genotype; control 11.21%±2.7, *Robo βKO* 25.99%2.9, *p*=0.02), and heterotypic contacts in general when compared to control islets (*n*=9-11 islets for 3 mice from each genotype; control 16.3%,±1.7 *Robo βKO* 35.57%±1.2, *p*=0.0008). Together, this suggests that *Robo βKO* islets make fewer homotypic β cell-β cell connections compared to control islets.

**Figure 1:**
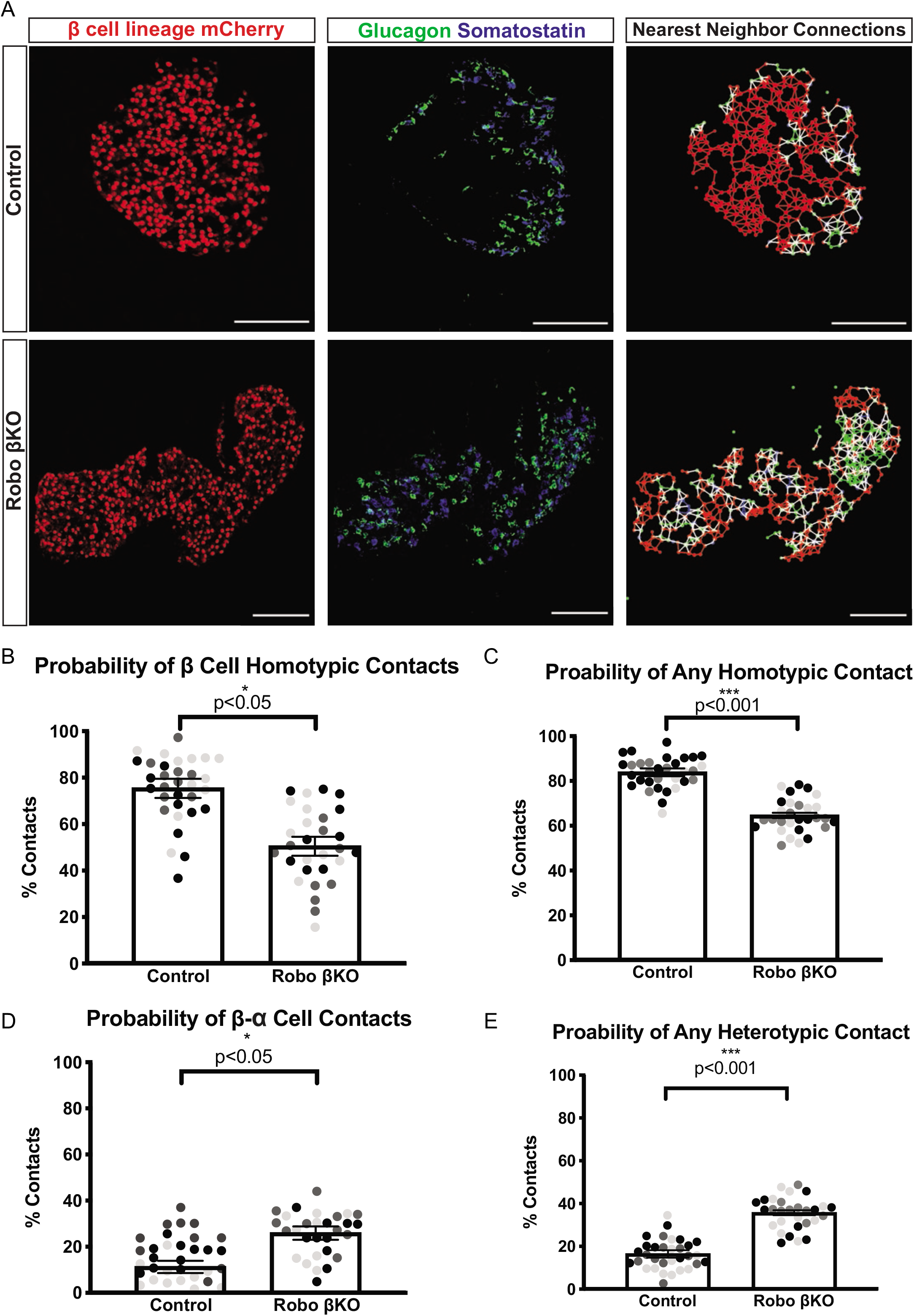
*Robo βKO* islets have a smaller fraction of homotypic nearest neighbors than controls. (A) Immunofluorescence images (left and middle panels) and cell connectivity maps generated by nearest neighbor analysis (right panels) of control and *Robo βKO* islets. β cells (red), α cells (green), and δ cells (blue) are denoted by nodes on the connectivity maps. A line the same color as both nodes it connects denotes a homotypic interaction of that corresponding cell type. A white line connecting two nodes denotes a heterotypic interaction between cell types. (B) Probability of β cell-β cell contacts in *Robo βKO* islets *vs*. controls (*n*=9-11 islets for 3 mice from each genotype; control 75.35%±4.1, *Robo βKO* 50.37%4.1, *p*=0.01). (C) Probability of any homotypic cell-cell contact in *Robo βKO* islets vs controls (*n*=9-11 islets for 3 mice from each genotype; control 83.7%±1.7, *Robo βKO* 64.43%±1.2, *p* 0.0008). (D Probability of β cell-α cell contacts in *Robo βKO* islets vs controls (*n*=9-11 islets for 3 mice from each genotype; control 11.21%±2.7, *Robo βKO* 25.99%±3.0, *p*=0.02). (E) Probability of any heterotypic cell-cell contact in *Robo βKO* islets *vs*. controls (*n*= 9-11 islets for 3 mice from each genotype; control 16.3%±1.7, *Robo βKO* 35.57%1.2, *p*=0.0008). (B-E Similar shaded points in graphs indicate islets from the same mouse).

We have previously shown that genetic deletion of *Robo1* and *Robo2* selectively in β cells using either *Ins1-Cre; Robo1^Δ/Δ^2^flx/flx^* or *Ucn3-Cre; Robo1^Δ/Δ^2^flx/flx^* mice (*Robo βKO*) results in disrupted islet architecture and endocrine cell type sorting without affecting β cell death or the expression of the β cell maturation markers MafA and Ucn3^36^. To verify that β cells in *Robo βKO* islet are truly mature, we expanded the analysis to look at transcript levels of 15 additional maturity markers. RNA sequencing and differential gene expression analysis on FACS-purified β cells from both *Robo βKO* and control islets revealed no change in transcript levels of any hallmark β cell maturity or differentiation genes (*n*=2 mice of each genotype; **Supplemental Figure 1**). Thus, unlike other mouse models with disrupted islet architecture, β cells in *Robo βKO* islets maintain maturity and differentiation despite loss of normal islet architecture.

We reasoned that the altered degree of homotypic β cell-β cell interaction in *Robo βKO* islets together with the seemingly retained β cell maturity provide a unique model by which to test the hypothesis that endocrine cell type organization affects synchronous insulin secretion in the islet.

### *Robo βKO* islets display unsynchronized Ca^2+^ oscillations *in vivo*

We set out to investigate how the reduced homotypic β cell-β cell connections in *Robo βKO* islets affects dynamic insulin secretion in the islet by measuring dynamic Ca^2+^ signaling. *Robo βKO* islets spontaneously dissociate during isolation and culture^36^, making them unsuitable for *in vitro* analyses of whole-islet Ca^2+^ oscillations. To overcome this limitation, we adopted a novel intravital Ca^2+^ imaging method which enables imaging of islet Ca^2+^ dynamics *in situ* within the intact pancreas^38^. In brief, this method employs an intravital microscopy (IVM) platform and adeno-associated viral (AAV) delivery of insulin promoter-driven GCaMP6s, a fluorescent Ca^2+^ biosensor, to quantitate β cell Ca^2+^ dynamics *in vivo* in both *Robo βKO* and control islets. This method also allows for retention of the islet’s *in vivo* microenvironment, blood flow, and innervation, and provides more realistic conditions than *in vitro* approaches allow for.

We verified that synchronous Ca^2+^ oscillations are maintained *in vivo* in islets by measuring GCaMP6s intensity of β cells within AAV8-RIP-GCaMP6 infected islets of control (*Robo WT*) mice (**Figure 2**). As expected, control mice displayed whole islet synchronous Ca^2+^ oscillations when imaged at 0.03, 0.1, and 0.2Hz for at least 10 minutes after glucose elevation (*n*=8 islets from 3 mice; **Figure 2** and **Supplemental Video 1 and 2**). We quantified the degree to which these oscillations are synchronous within the islet by analyzing the amount of correlation between GCaMP6s active areas within individual islets. While oscillations vary in frequency between islets, the degree of correlation between β cells within any one islet is very high, confirming that control islets possess highly synchronous intra-islet Ca^2+^ oscillation *in vivo* (fraction of GCAMP6s active islet area with correlated Ca^2+^ oscillations=0.90±0.04, *n*=8 islets from 3 mice; see **Figure 4A**).

**Figure 2:**
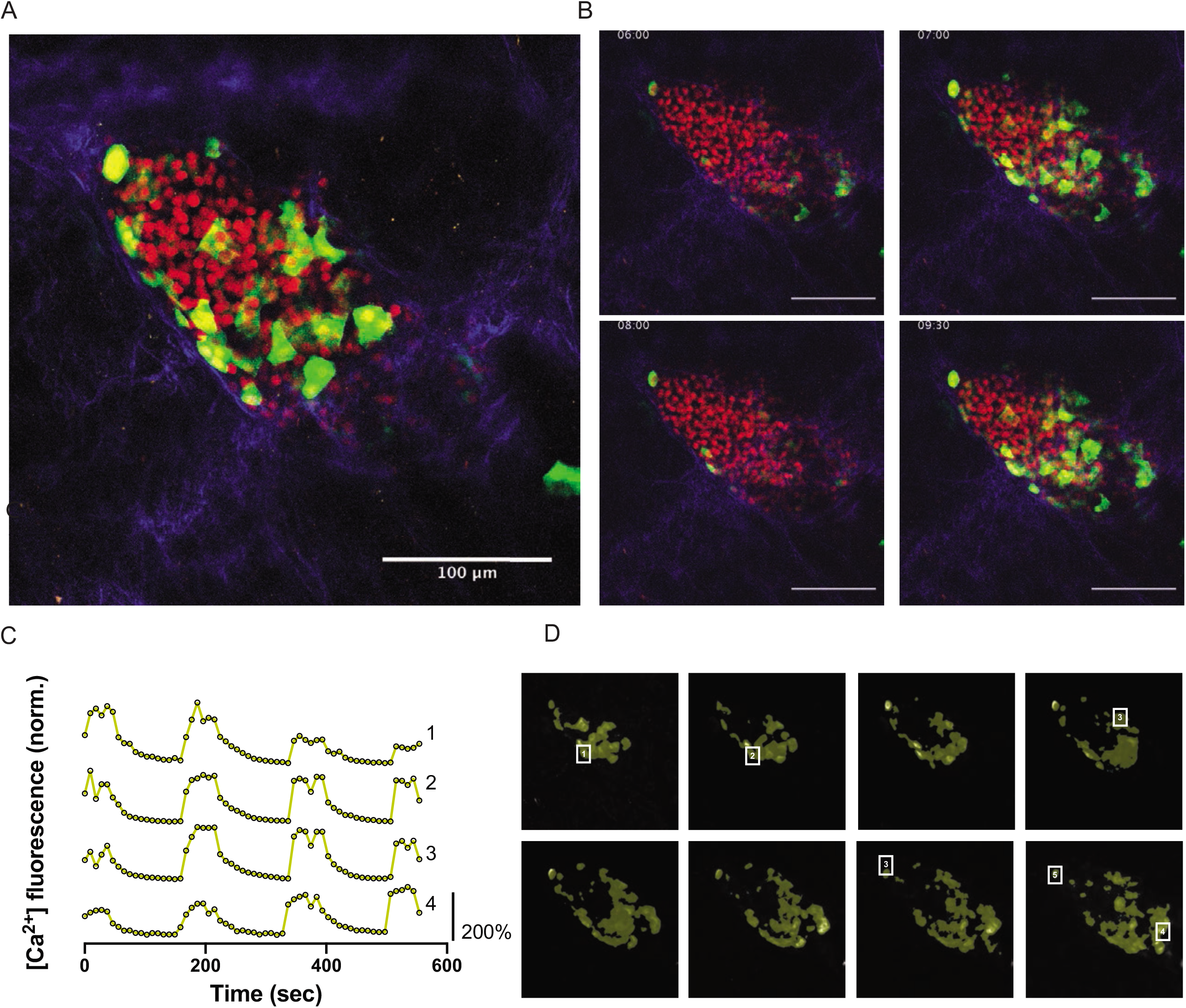
Control islets show highly synchronized whole islet Ca^2+^ oscillations. (A) High resolution maximum intensity projection of a control islet *in vivo* in an *AAV8-RIP-GCaMP6s*-injected mouse showing GCaMP6s in green, nuclear mCherry β cell lineage-tracing in red, and collagen (second-harmonic fluorescence) in blue. (B) Stills over one oscillation period from control islet in supplementary video 1, starting after blood glucose level reached ^~^300mg/dL from IP glucose injection. Video was recorded for 10 minutes with an acquisition speed of 0.1Hz. (C) Representative time courses of Ca^2+^ activity in 4 individual areas from control islet in supplementary video 1 showing correlation over 98% of the active islet area. Time courses are normalized to average fluorescence of individual area over time. Similar color indicates that the time courses have a Pearson’s correlation coefficient of ≥0.75 and matches the region of coordination that is seen in D. (D) False color map of top five largest coordinated areas across z-stack of control islet from analysis in C. Areas in grey are not coordinated. The color represents a region of coordination with Pearson’s Correlation Coefficient ≥0.75 of GCaMP6s activity. Cells used in time courses in C are labeled.

Conversely, we found that most *Robo βKO* islets display asynchronous intra-islet Ca^2+^ oscillations *in vivo* when imaged at 0.03, 0.1, and 0.2Hz (**Figure 3, Supplemental Figure 2, Supplemental Videos 3, 4, and 5**). Quantification of this asynchronous behavior through correlation analysis of GCaMP6s activity within individual *Robo βKO* islets revealed significant reduction in intra-islet correlated oscillation areas compared to controls (fraction of GCAMP6s activity with correlated Ca^2+^ oscillations=0.58±0.10, *n*= 11 islets from 5 mice, *p*<0.01; **Figure 4A**). Further, asynchronous *Robo βKO* islets showed spatially distinct areas within the islet that oscillated synchronously with immediate β cell neighbors but not with more distant regions within the same islet (**Figure 3C-D, Supplemental Figure 2, Supplemental Videos 3, 4, and 5**). This was not due to differences in the proportion of GCaMP6s positive cells showing elevated Ca^2+^ activity within *Robo βKO* islets compared to controls (control islets 0.97±0.01 fraction active, *n*=8 islets from 3 mouse; *Robo βKO* islets 0.97±0.02 fraction active, *n*=8 islets from 4 mice, *p*=0.79; **Figure 4B**).

**Figure 3:**
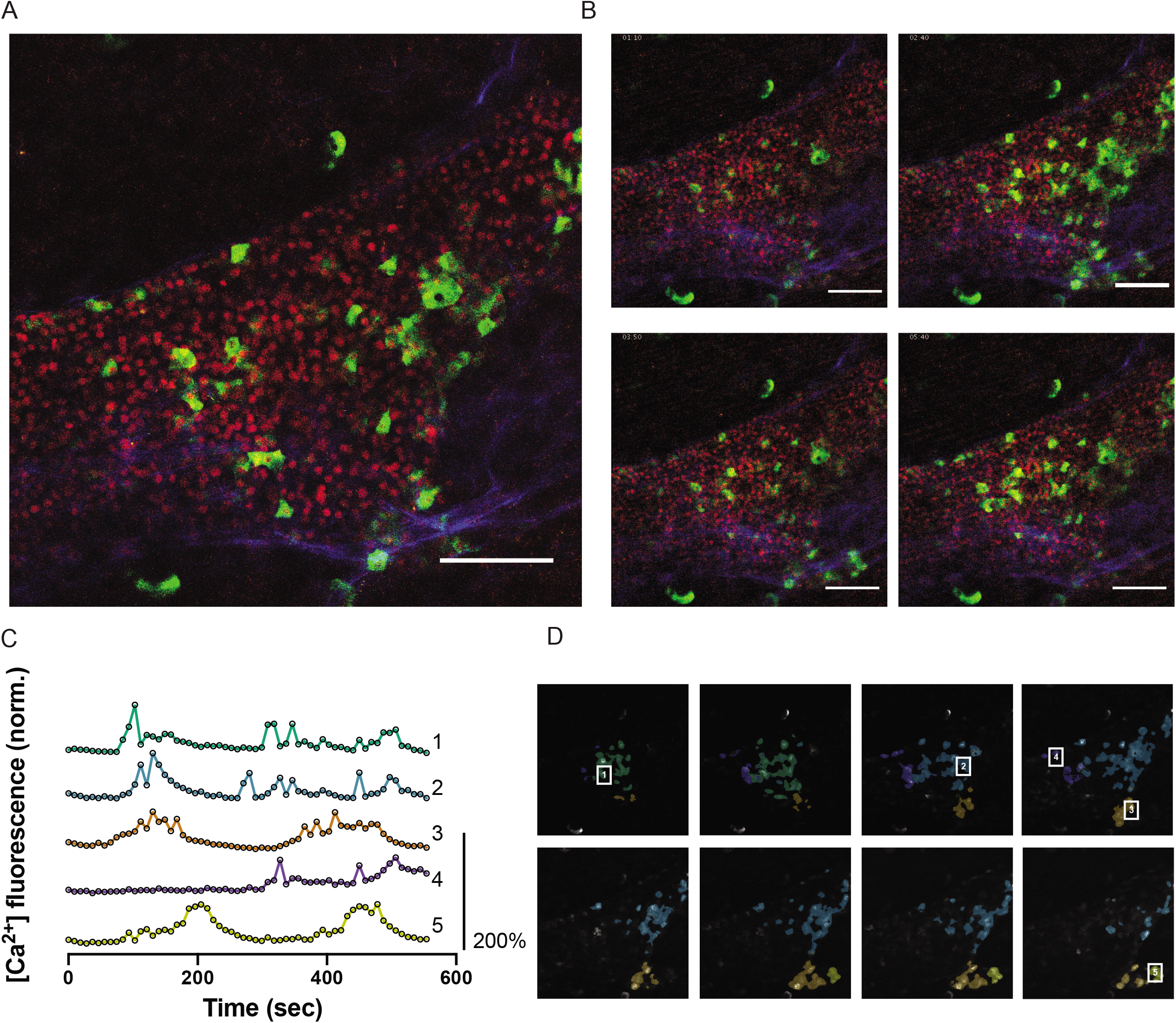
*Robo βKO* islets show uncoordinated whole islet Ca^2+^ oscillations. (A) High resolution maximum intensity projection of a *Robo βKO* islet *in vivo* in an *AAV8-RIP-GCaMP6s-injected* mouse showing GCaMP6s in green, nuclear mCherry β cell lineage-tracing in red, and collagen (second-harmonic fluorescence) in blue. (B) Stills over one oscillation period from *Robo βKO* islet in supplemental video 4, starting after blood glucose level reached ^~^300mg/dL from IP glucose injection. Video was recorded for 10 minutes with an acquisition speed of 0.1Hz. (C) Representative time courses of Ca^2+^ activity in 4 individual areas from *Robo βKO* islet in supplementary video 4, showing correlation of 50% of the active islet area. Time courses are normalized to average fluorescence of individual area over time. Similar color indicates that the time courses. Similar color indicates that the time courses have a Pearson’s correlation coefficient of ≥0.75 and matches the region of coordination that is seen in D. (D) False color map of top five largest coordinated areas across z-stack of *Robo βKO* islet from analysis in C. Areas in grey are not coordinated. The color represents a region of coordination with Pearson’s correlation coefficient of ≥0.75 of GCaMP6s activity.

**Figure 4:**
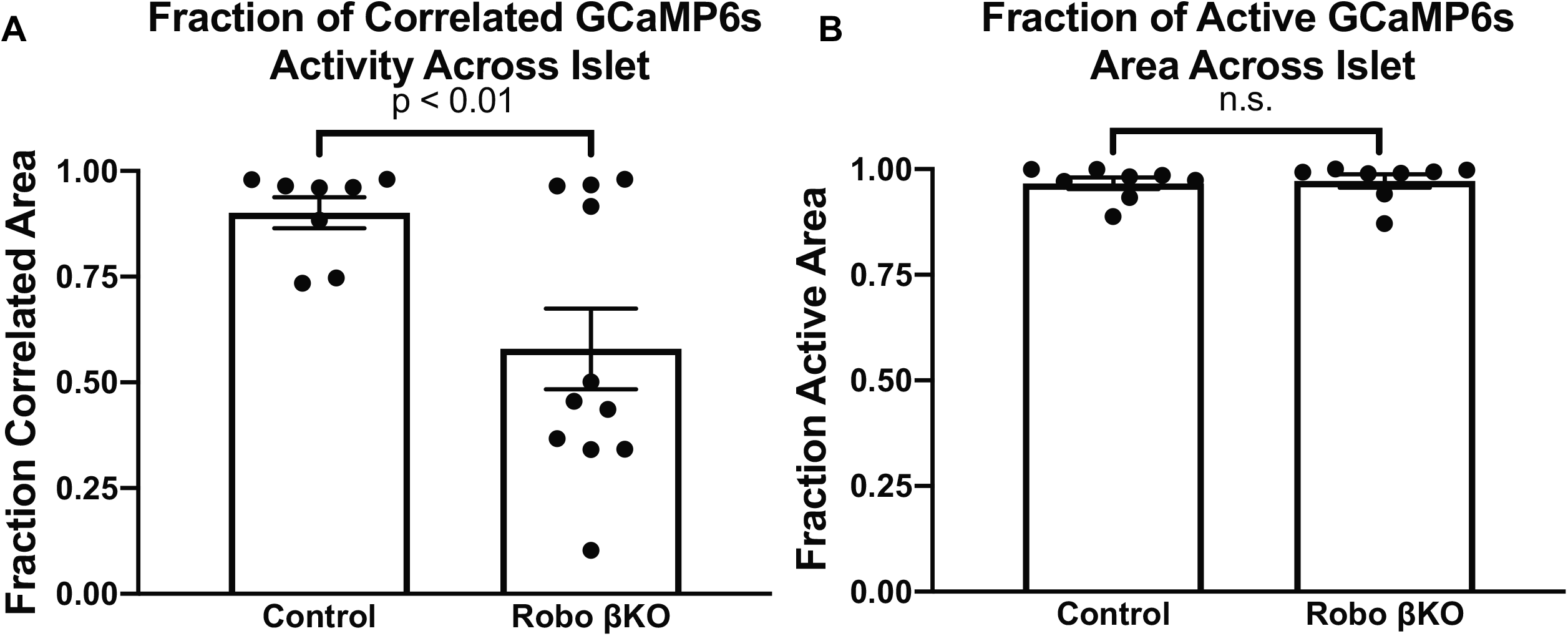
Quantification of *Robo βKO* Ca^2+^ oscillation phenotype. (A) Largest fraction of area in islet exhibiting coordinated Ca^2+^ oscillations for control and *Robo βKO* islets. (B) Fraction of active islet area showing elevated Ca^2+^ activity for control and *Robo βKO* islets.

### Unsynchronized Ca^2+^ oscillations in Robo βKO islets are not due to β cell intrinsic defects in glucose stimulated Ca^2+^ oscillations

*In vitro* experiments have shown that Robo receptors play a role in β cell biology and are involved in the stimulus secretion cascade linking glucose to insulin secretion^39^. Thus it is possible that defects in synchronous Ca^2+^ oscillations observed in *Robo βKO* islets are due to a Robo-mediated, β cell-intrinsic effect on dynamic Ca^2+^ signaling in response to stimuli. However, 4 out of the 11 *Robo βKO* islets imaged showed synchronous Ca^2+^ activity in greater than 90% of GCaMP6s positive areas (**Figure 4A, Supplemental Figure 3 and Supplemental video 6**). The existence of this highly synchronous population of *Robo βKO* islets suggests that the ability of individual β cells to oscillate intracellular Ca^2+^ levels in response to stimuli is unaffected by deletion of *Robo*. Instead, it suggests that islet architecture itself is responsible for controlling synchronized Ca^2+^ oscillations, and that some *Robo βKO* islets may escape this defect due to less severe architectural disruption. With this is mind, we wanted to test the extent to which the unsynchronized Ca^2+^ oscillations observed in *Robo βKO* islets *in vivo* are due to β cell-intrinsic deletion of *Robo per se*.

To test whether *Robo βKO* β cells are able to undergo Ca^2+^ oscillations in response to stimuli, we performed *in vitro* Ca^2+^ imaging on single β cells from dissociated *Robo βKO* and control islets, exposed to glucose followed by KCL (**Figure 5**). We found no difference in the proportion of β cells that undergo Ca^2+^ oscillations in response to 10mM glucose between control and *Robo βKO* β cells (Control: 70.24%±3.6, *n*=13-32 β cells per mouse for 4 mice, *Robo βKO*: 61.61%±5.6, *n*=6-24 β cells per mouse for 4 mice, *p*=0.24) (**Figure 5 A-C**). We also saw no significant difference in area under the curve (AUC) of calcium traces in response to 10mM glucose (Control: 2.9±0.17, *n*=13-32 β cells per mouse for 4 mice, *Robo βKO:* 3.0±0.21, *n*=6-24 β cells per mouse for 4 mice, *p*=0.39)(**Figure 5D**), or peak height of Ca^2+^ corresponding to first phase insulin secretion (Control: 0.34±0.02, *n*=13-32 β cells per mouse for 4 mice, *Robo βKO:* 0.41±0.04, *n*=6-24 β cells per mouse for 4 mice, *p*=0.08) (**Figure 5E**) in *Robo βKO* β cells compared to controls. There was, however, a significant increase in AUC of Ca^2+^ signal in response to KCL in *Robo βKO* β cells compared to controls (Control: 1.67±0.07, *n*=13-32 β cells per mouse for 4 mice, *Robo βKO*: 2.12±0.12, *n*=6-24 β cells per mouse for 4 mice, *p*=0.002) (**Figure 5F**), indicating that *Robo βKO* islets may actually have an increase in magnitude of Ca^2+^ response to stimuli. Together, this demonstrates that *Robo βKO* β cells show no defects in their ability to undergo Ca^2+^ oscillations in response to glucose, and in fact may have an improved stimulus secretion response compared to controls. This is in support of a scenario in which the arrangement of β cells within the islet, rather than a β cell-intrinsic function of Robo, dictates whole-islet synchronous Ca^2+^ oscillations.

**Figure 5:**
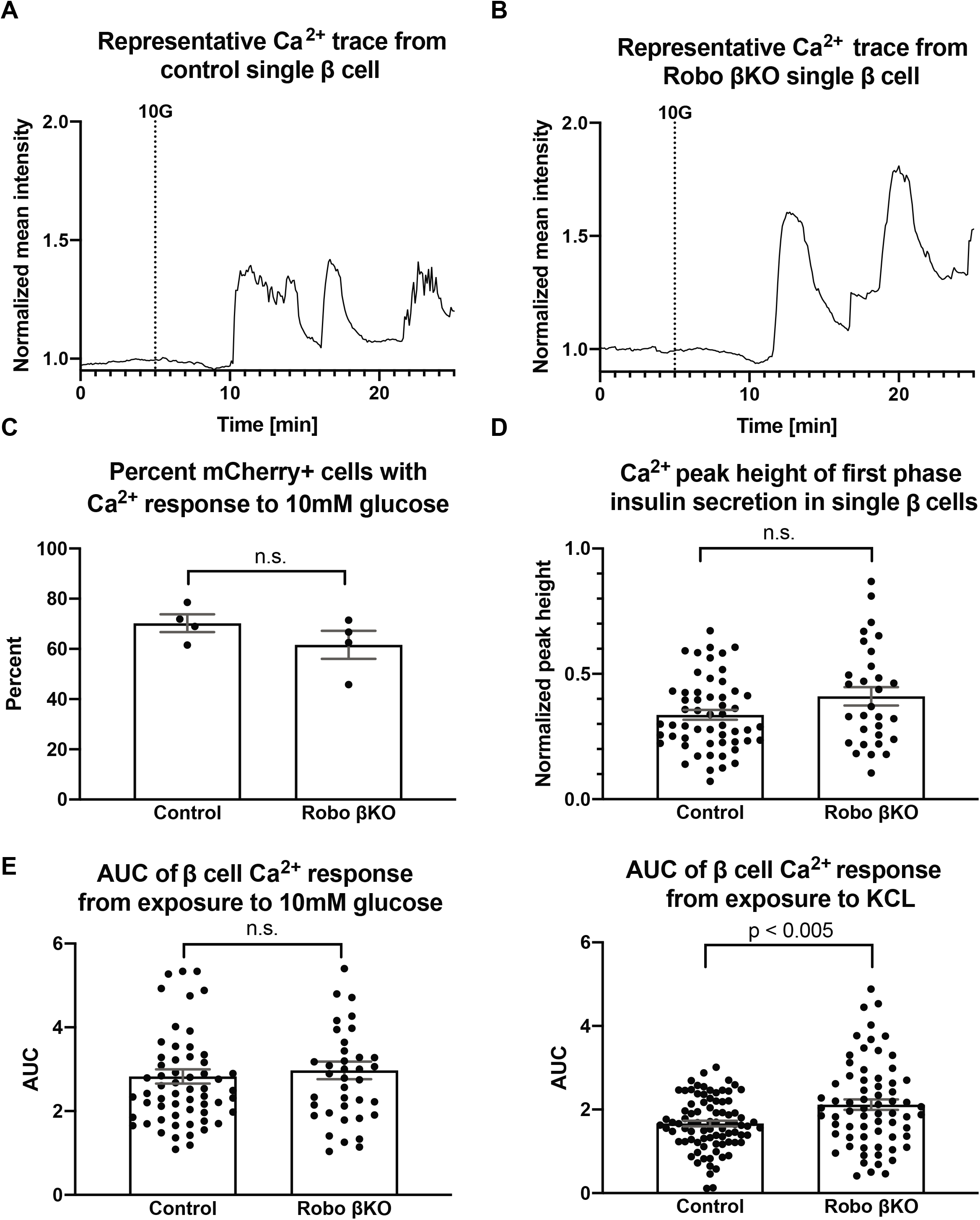
Dissociated *Robo βKO* β cells show no difference in glucose stimulated Ca^2+^ oscillations. (A) Representative Ca^2+^ trace (Fura2) of a single dispersed β cell from a control islet. 10G line marks the addition of 10mM glucose. (B) Representative Ca^2+^ trace (Fura2) of a dispersed β cell from a *Robo βKO* islet. 10G line marks the addition of 10mM glucose. (C) Graph showing the proportion of Ca^2+^ responsive β cells in *Robo βKO* compared to controls (D) Graph showing peak height of Ca^2+^ oscillation corresponding to first phase insulin secretion from control and *Robo βKO* single dispersed β cells in response to 10mM glucose. (E) Graph showing area under the curve (AUC) of Ca^2+^ oscillations (Fura2) from control and *Robo βKO* single dispersed β cells in response to 10mM glucose. (F) Graph showing area under the curve (AUC) of Ca^2+^ oscillations (Fura2) from control and *Robo βKO* single dispersed β cells in response to KCL.

### *Robo βKO* islets retain the ability to form gap junctions and have similar levels of vascularization

Besides a decrease in β cell-β cell homotypic contacts within the islet, a possible explanation for the loss of synchronized whole islet Ca^2+^ oscillations in *Robo βKO* is that their β cells no longer possess the gap junctions necessary for adequate electrical coupling. Indeed, the phenotype described above is reminiscent of that observed in mice heterozygous for a *Cx36* null allele^15,40^. To test whether *Robo βKO* mice form fewer gap junctions between β cells, we measured the area of Cx36 protein immunofluorescence normalized to islet area in *Robo βKO* and control islets (**Figure 6**). We found no difference in Cx36 immunofluorescence between *Robo βKO* islets and controls, but significant differences in these two groups compared to *Cx36 KO* mice (control islets: 0.46%±0.06 Cx36 signal/μm^2^, *Robo βKO* islets: 0.69%±0.11 Cx36 signal/μm^2^, *n*=10-15 islets from 4 mice each, p=0.11, *Cx36 KO* islets: 0.13%±0.04, n=15-20 islets from 2 mice) (**Figure 6B**). Furthermore, we found normal co-localization of Cx36 to β cell borders in both control and *Robo βKO* islets (**Figure 6C**) Overall, this suggests that loss of synchronous intra-islet Ca^2+^ oscillations is not due to failure of gap junction formation in β cells.

**Figure 6:**
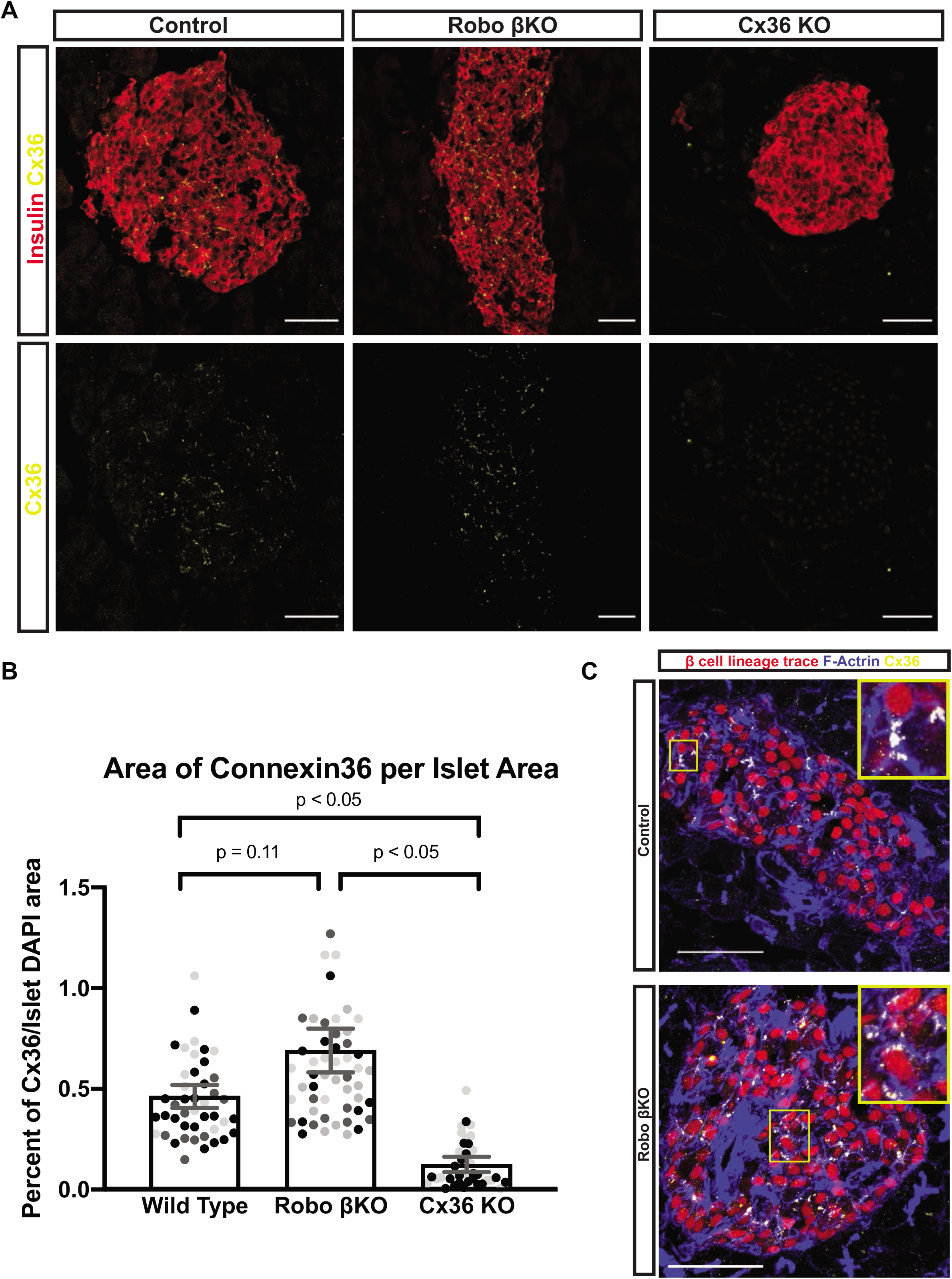
Amount of Cx36 gap junctions remains unchanged in *Robo βKO*. (A) Immunofluorescent images showing Cx36 (gray or green) and insulin (red) in Control, *Robo βKO*, and *Cx36 KO* islets. (B) Quantification of area of Cx36 staining normalized to islet area in *Robo βKO* islets and controls showing no significant difference (*n*=10-20 islets from 2-4 mice per group, *p values shown*). similar colored dots represent islet from the same mouse (C) Immunofluorescent images showing histone H2B-mCherry β cell lineage trace in red, F-Actin (phalloidin) in blue, and Cx36 in yellow demonstrating normal localization of Cx36 to plasma membrane (visible as white dots) of β cells in both control and *Robo βKO* islets.

Another possible explanation for the observed uncoupling of intra-islet Ca^2+^ oscillations in *Robo βKO* islets is that β cell-β cell contacts are disrupted due to physical blocking by non-endocrine tissue. To determine if other non-endocrine architectural changes within the islet occur in *Robo βKO* mice we quantified the amount of matrix components secreted by vessels as a surrogate for vasculature (laminin, and collagen IV) in *Robo βKO* and control islets. In both cases we found no significant difference in area of vessel matrix components (**Figure 7**) between *Robo βKO* and control islets, suggesting that interfering blood vessels are likely not the cause of loss of whole islet synchronous Ca^2+^ oscillations (normalized laminin β1 area: control 0.31±0.05 μm^2^, *n*=10-12 islets from 8 mice; *Robo βKO* 0.38±0.05 μm^2^, *n*=10-12 islets from 8 mice; *p*= 0.39; normalized Col IV area: control 0.18±0.01 μm^2^, *n*=10-12 islets from 4 mice; *Robo βKO* 0.16±0.02 μm^2^, *n*=10-12 islets from 6 mice, *p*=0.35).

**Figure 7:**
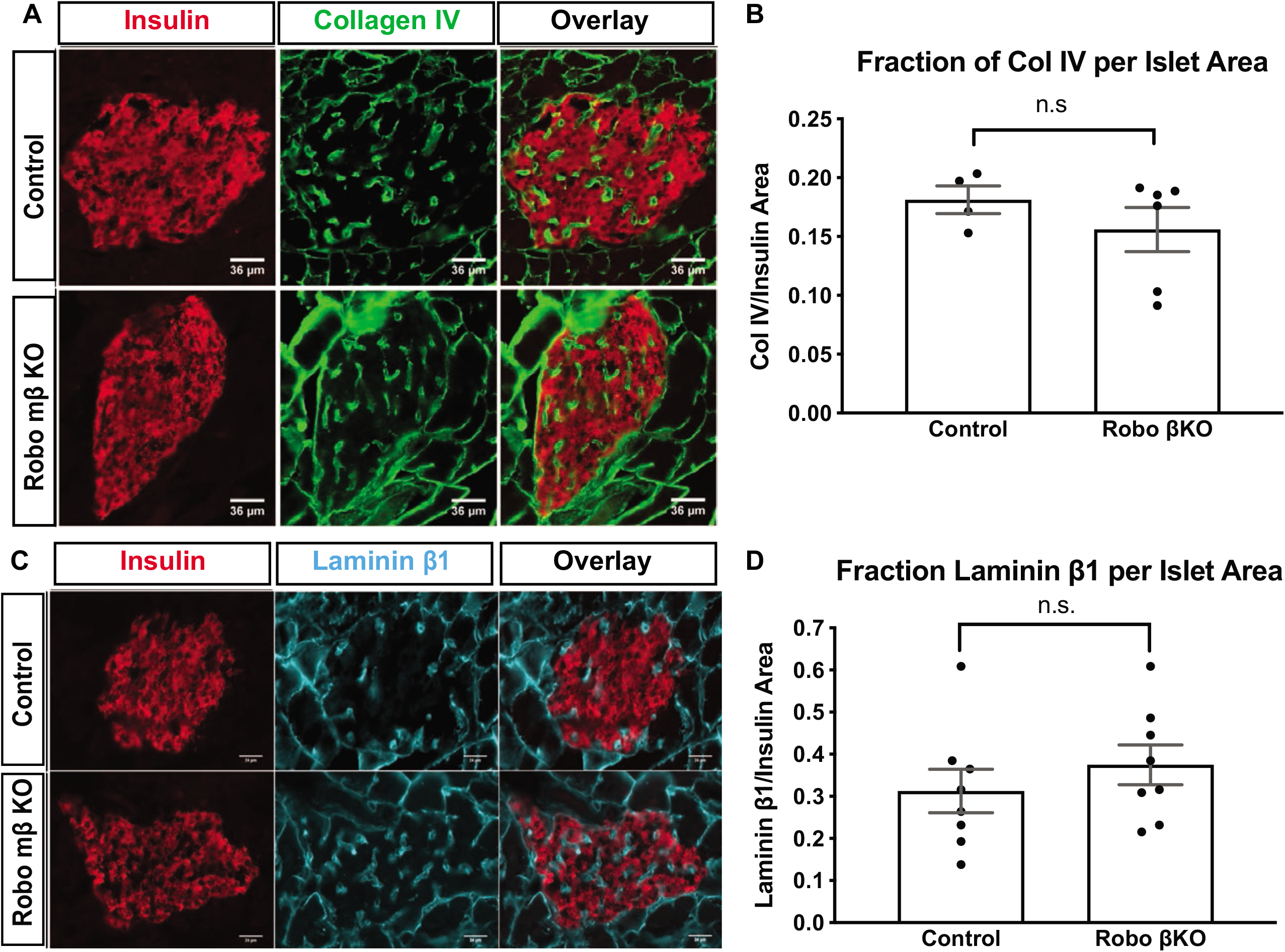
Amount of vascularization remains unchanged in *Robo βKO* islets. (A) Representative immunofluorescent staining of Collagen IV marking vasculature showing similar amounts in *Robo βKO* and control islets (B) Quantification of area of Collagen IV staining normalized to islet area showing no difference in amounts of basement membrane marking blood vessels in *Robo βKO* compared to control islets. (C) Representative immunofluorescent staining of laminin marking vasculature showing similar amounts in *Robo βKO* and control islets (D) Quantification of area of laminin staining normalized to islet area showing no difference in amounts of basement membrane marking blood vessels in *Robo βKO* compared to control islets.

## Discussion

In this study, we provide evidence for the importance of islet architecture for proper islet function *in vivo*. When islet architecture is disrupted while β cell maturity is retained in *Robo βKO* mice, synchronized Ca^2+^ oscillations in the islet *in vivo* are perturbed. This is not due to a Robo-mediated β cell-intrinsic defect in glucose-stimulated Ca^2+^ signaling, loss of Cx36, or change in amount of islet vascularization. *Robo βKO* β cells possess a smaller fraction of homotypic nearest neighbors than controls, suggesting a limited capacity to electrically couple β cells across the islet. Taken together, these data suggest that islet architecture itself, uncoupled from β cell maturity, Robo-mediated β cell-intrinsic defects in Ca^2+^ signaling, or availability of gap junction machinery, is important for coordinated insulin secretion between β cells.

Robo, and its ligand Slit, have been previously shown to affect Ca^2+^ oscillations in β cells *in vitro*^39^. However, Robo-mediated β cell-intrinsic effects are likely not the cause of asynchronous *in vivo* Ca^2+^ oscillations in *Robo βKO* islets. This is supported by two independent observations: 1) single cell Ca^2+^ oscillations in dissociated β cells triggered by glucose stimulus *in vitro* are similar between *Robo βKO* and control, and 2) a subset of *Robo βKO* islets analyzed *in vivo* still show synchronized whole-islet Ca^2+^ oscillations despite the absence of Robo. Further, this phenotypic heterogeneity is likely not due to an incomplete deletion of *Robo* because all islets were detected during the Ca^2+^ imaging experiment by the fluorescent labeling of β cells with the H2B-mCherry lineage-tracing reporter^36^ which uses the same Cre deriver that is used to delete *Robo* in those β cells. Thus high expression of H2B-mCherry suggests efficient recombination of the *Robo* floxed allele. All this together supports the idea that *Robo* deletion in β cells causes disruption of islet architecture, and that this architectural disruption itself causes loss of synchronized Ca^2+^ oscillations. Further, the unsynchronized phenotype in *Robo βKO* islets is clear from the videos we captured at 0.2Hz, 0.1Hz, and 0.03Hz; however we are aware that even higher imaging frequency may be needed to see asynchronous behavior in *Robo WT* islets. That said, we note that while faster acquisition would be beneficial to show more rapid dynamics, we believe it important to capture the dynamics in 3D, in order to ensure we are not biased by examining certain ‘synchronized regions’ that would not be representative of the islet dynamics as a whole. We would further argue that the oscillations we observe are on a 3-5 minute time scale, and therefore to sample these oscillations even at a frequency of 1 frame per 10s of seconds is sufficient. Indeed if we were to image more rapidly and capture other non-synchronized dynamics in the *Robo βKO* islets, this would reduce the measured coordination even further. Thus the low sample frequency provides an upper limit to the coordination.

It remains possible that other components of islet architecture besides endocrine cell type sorting contribute to disruption in Ca^2+^ oscillations found in *Robo βKO* islets. While we have shown that the amount of vascularization between *Robo βKO* islets and controls is similar, we cannot draw conclusions on whether the pattern of vessels is unchanged. Further, it is possible that the amount and patterning of innervation may vary between *Robo βKO* and controls. These are particularly of interest because Robo receptors have known roles in angiogenesis and axon guidance, and thus could affect precisely how the islet is innervated and vascularized^41^. Robo also has known roles in controlling cell polarity, and thus it is possible that this process is disrupted in *Robo βKO* islets as well. Gap junctions are known to be localized to the junctional membranes between β cells, which themselves are defined by β cell polarity within the islet^40,42^. Thus if β cell polarity is disrupted in *Robo βKO* islets, this could possibly contribute to the asynchronous Ca^2+^ oscillations observed.

Finally, while we propose that defects in synchronous Ca^2+^ oscillation in response to glucose in *Robo βKO* islets are due to inefficient electrical β cell coupling as a result of decreased homotypic β cell interactions in these islets^43^, we cannot rule out that this effect is instead due to changes in diffusible paracrine factors. Under this hypothesis, the intermixed islet architecture that results from deletion of *Robo* in β cells would change the amount and type of diffusible paracrine signals that endocrine cells within the islet are exposed to. This change in diffusible factors could affect glucose stimulated insulin secretion and thus could be contributing to the *in vivo* asynchronous Ca^2+^ oscillation phenotype that we observe^44^.

## Acknowledgments

We thank Kurt Weiss, Jan Huisken and David Inman for help with imaging. We thank members of the Blum lab, especially Jennifer Gilbert and Bayley Waters, for valuable discussion. We are also grateful to Nadav Sharon and Danny Ben-Zvi for critically reading the manuscript. This work was funded in part by the following grants. R01DK121706, the DRC at Washington University Pilot Grant P30DK020579, and Pilot Award UL1TR000427 from the UW-Madison Institute for Clinical and Translational Research (ICTR) to BB; R01DK060581 to RM; R01DK102950 and R01DK106412 to RKPB; R01CA216248 to SP; R01DK113103, R01AG062328, ADA 1-16-IBS-212 to MM, and an award from the Wisconsin Partnership Program to BB and MM. MTA was funded by 5T32GM007133-44 and a graduate training award from the UW-Madison Stem Cell & Regenerative Medicine Center. We also thank the University of Wisconsin Carbone Cancer Center Support Grant P30CA014520 for use of the UW Flow Core, and the University of Wisconsin, Madison Biotechnology Center for sequencing and analysis.

## Author Contribution

Conceptualization, B.B. and M.T.A; Methodology, B.B., M.T.A., C.A.R, and J.M.D.; Investigation, M.T.A., C.A.R, J.M.S, M.R.L, S.M.S,, E.J, and S.D.N; Formal Analysis, M.T.A., C.A.R, M.R.L., E.J, and J.M.D.; Resources, S.M.P, R.G.M, A.K.L, M.J.M. and R.K.P.B.; Writing Original Draft, B.B and M.T.A.; Writing, Review and Editing, all authors; Funding Acquisition, B.B., S.M.P, M.J.M, R.K.P.B., R.G.M., and A.K.L; Supervision, B.B.

## Materials and Methods

### Animals

The experimental protocol for animal usage was reviewed and approved by the University of Wisconsin-Madison Institutional Animal Care and Use Committee (IACUC) under Protocol #M005221 and Protocol #M005333, and all animal experiments were conducted in accordance with the University of Wisconsin-Madison IACUC guidelines under the approved protocol. *Robo1*^Δ^,2^*flx*45^, *Ins1-Cre*^46^, *Urocortin3-Cre*^47^ and *Rosa26-Lox-Stop-Lox-H2BmCherry*^48^ mice were previously described. All mouse strains were maintained on a mixed genetic background. Control colony mates in all analyses were *Robo^+/+^* with the either *Ins1-Cre or Ucn3-Cre*.

### Immunofluorescence

Pancreata were fixed with 4% PFA at 4°C for 3h, embedded in 30% sucrose and frozen in OCT (Tissue-Tek). Pancreatic sections (10 μm) were stained using a standard protocol. The following primary antibodies and dilutions were used: guinea pig anti-Insulin (1:6, Dako, IR00261-2), mouse anti-Glucagon (1:500, Sigma G2654), rabbit anti-Glucagon (1:200, Cell Signaling 2760S), rabbit anti-Somatostatin (1:1000, Phoenix G-060-03), rabbit anti-Connexin36 (1:80, Invitrogen 36-4600), rabbit anti-Col IV (1:300, Abcam Ab656), rat anti-Laminin β1 (1:500, Invitrogen MA5-14657). The following secondary antibodies were used at 1:500: Donkey anti-Guinea Pig 594 (Jackson), Donkey anti-Guinea Pig 647 (Jackson), Donkey anti-Rabbit 488 (Invitrogen), Donkey anti-Rabbit 594 (Invitrogen), Donkey anti-goat 647 (Invitrogen), and Donkey anti-rat 488 (Invitrogen). Slides were imaged using a Leica SP8 Scanning Confocal microscope or a Zeiss Axio Observer.Z1 microscope.

### RNA sequencing

RNA was isolated from FACS sorted lineage-traced β cells^36^ from control and *Robo βKO* mice using phenol chloroform extraction (TRIzol). DNA libraries were generated using Takara’s SMART-Seq v4 Low Input RNA Kit for Sequencing (Takara, Mountain View, California, USA) for cDNA synthesis and the Illumina NexteraXT DNA Library Preparation (Illumina, San Diego, CA, USA) kit for cDNA dual indexing. Full length cDNA fragments were generated from 1-10ng total RNA by SMART (Switching Mechanism at 5’ End of RNA Template) technology. cDNA fragments were fragmented and dual indexed in a single step using the Nextera kit’s simultaneous transposon and tagmentation step. Quality and quantity of completed libraries were assessed using Agilent DNA series chip assay (Agilent Technologies, Santa Clara, CA) and Invitrogen Qubit ds DNA HS Kit (Invitrogen, Carlsbad, California, USA), respectively. Each library was standardized to 2nM. Cluster generation was performed on Illumina cBot, with libraries multiplexed for 1×100bp sequencing using TruSeq 100bp SBS kit (v4) on an Illumina HiSeq2500. Images were analyzed using standard Illumina Pipeline, version 1.8.2.

### Intravital imaging

Mouse pancreata were exposed in anesthetized mice by making a small incision on the right side of the mouse, and externalizing the tip of the pancreas. A glass dish was placed over the exposed pancreas and the mouse was placed on a microscope stage with isoflurane anesthesia for the remainder of imaging. Islets were identified on the surface of the pancreas by detecting Histone H2B-mCherry fluorescent nuclei labeled by β cell-specific lineage-tracing reporter^36^. Once islets were identified, mice were given injections of 1g/kg body weight glucose (30% in saline) intraperitoneally. Blood glucose levels were monitored through tail vein bleeds. Once the blood glucose reached ^~^300 mg/dL, GCaMP6s activity was identified using the microscope eye piece. When imaging a time course of GCaMP6s intensity, a z-stack was set to 3, 8 or 12 slices each 8μm apart. Images were captured at 0.2Hz, 0.1Hz, or 0.03Hz respectively over at least 10 minutes at a resolution of 512×512 pixels. After time courses were recorded, high resolution image z-stacks were taken with 60 z planes taken 1μm apart or 8 z-planes taken 8μm apart at 1024×1024 pixel resolution. For some images, rhodamine-dextran was injected retro-orbitally to mark the vasculature of the islets *in vivo*.

### Gap junction and vasculature quantification

Cx36 levels were quantified from images of islets co-stained with rabbit anti-Cx36 (Invitrogen) and Guinea Pig anti-insulin (Dako) antibody. Vasculature levels were quantified from images co-stained with rat anti-Laminin β1 (Invitrogen) or rabbit anti-col IV and Guinea Pig anti-insulin (Dako). 8 Z-planes were taken 1μm apart on a Leica SP8 Scanning Confocal microscope using a 40x oil immersion objective (Cx36) or 20X (vasculature). Threshold masks were made of both channels for each islets, and the area of each staining was measured using FIJI’s analyze particles functions. The area of gap junctions or blood vessels (marked by their respective antibody) was divided by the area of DAPI or insulin respectively for each islet. 10-14 islets were analyzed for *n*=3-8 mice for each genotype. Student’s T-test was performed to obtain *P* values.

### Nearest neighbor analysis

β cells were identified using the lineage tracer *Rosa26-Lox-Stop-Lox-H2BmCherry* crossed to *Ucn3-Cre* and tissue sections were stained with antibodies against glucagon and somatostatin to identify α and δ cells respectively. The 3D Tissue Spatial Analysis Toolbox for Fiji^49^ was used to identify specific cell types using the above markers and to calculate the number of cell type specific nearest neighbors from all identified endocrine cells. Analysis was performed on 9-11 islets from *n*=3 mice from each genotype.

### Time course image analysis

All images were analyzed using previously published methods^50^ with custom Matlab (Mathworks) scripts. For activity analysis, images were smoothed using a 5×5 pixel averaging filter. Areas without significant fluorescence were removed. Saturated areas were also removed by limiting the area to intensity below the maximum value. Photobleaching was adjusted for by removing any linear trend. Any islets with significant motion artifacts were removed or time courses were shortened to the time over which no significant movement occurred (displacement of <0.5 cell width). For the time course of each pixel in the image with significant fluorescence, a peak detection algorithm was used to determine if the areas had peak amplitudes significantly above background. A region was considered “active” if the corresponding time course for each pixel had a peak amplitude >2.4x background. The fraction of active area was calculated as the number of pixels detected as “active” across all z-planes, normalized to the total number of pixels that showed significant fluorescence across all z-planes that were not saturated. Islets with significant background fluorescence from spectral overlap of channels were excluded from activity analysis because “inactive” cells were indistinguishable from background and therefore total islet area could not be accurately calculated. Coordination was determined based on coincident timing of identified peaks, where areas were segmented by identified peaks occurring at similar time points. The cross correlation of the time courses for two 5×5 pixel subregion was taken. If the correlation coefficient was >0.75, then the two subregions were considered highly coordinated and merged into a larger region. The coordinated area was calculated as the number of pixels in the largest area of coordination across all z-planes normalized to the total number of pixels of the islet that were determined to be ‘active’ for all planes. This analysis is based on previous analysis^50^, but adjusted for 3-dimensional data. All statistical analysis was performed in Prism (Graphpad) or Matlab. First a F-test was used to determine if variances were equal then a Student’s t-test or Welch t-test (for unequal variance) were utilized for determining whether activity, coordination, phase lag and speed were significantly different. *p*<0.05 was considered significant.

### *In vitro* single cell Ca^2+^ imaging

Islets were isolated according to standard protocol from 3-6 month old *Robo βKO* and control mice. For islet dispersion, 12mm round No. 1.5 coverslips contained in a 24-well plate were pre-coated overnight with 50μL 1:15000 PEI (Sigma P3143) overnight. Groups of 100 mouse islets were dispersed into single cells in 3mL Accutase (Thermo Fisher A1110501) at 37°C for 10 min. During the incubation, PEI was replaced with 100μL Geltrex (Thermo Fisher A1413302) and centrifuged at 500*g* for 5 min at 4°C, followed by removal of excess Geltrex. The cells were washed once with islet culture medium (RPMI1640 supplemented with 10% FBS (v/v), 100 units/mL penicillin, and 100ug/mL streptomycin (Invitrogen)) and resuspended in 1mL medium before plating 500μL per coverslip. The plate was centrifuged for 5 minutes at 500*g* and cultured overnight before imaging. For measurements of cytosolic Ca^2+^, dispersed islet cells were pre-incubated in 5μM Fura2-AM (Thermofisher F1201) in islet media containing 11.1mM glucose for 45 min at 37°C, followed by 15 min incubation in islet media containing 2.7mM glucose. Coverslips were transferred to a RC-48LP imaging chamber (Warner Instruments) mounted on a Nikon Ti-Eclipse inverted microscope equipped with a 20X/0.75N.A. SuperFluor objective and PerfectFocus (Nikon Instruments). The chamber was perfused with a standard external solution containing 135mM NaCl, 4.8mM KCl, 2.5mM CaCl_2_, 1.2mM MgCl_2_, 20mM HEPES, and glucose as indicated (pH 7.35). The flow rate was set to 0.4mL/min (Fluigent MCFS-EZ) and temperature was maintained at 33°C using solution and chamber heaters (Warner Instruments). Excitation was provided by a SOLA SE II 365 (Lumencor) set to 10% output and an inline neutral density filter (Nikon ND4). Fluorescence emission was collected with a Hamamatsu ORCA-Flash4.0 V2 Digital CMOS camera at 0.1Hz. Excitation (x) and emission (m) filters were used in combination with a ET FURA2/GFP C164605 dichroic (Chroma): Fura2, ET365/20x, ET535/30m; mCherry ET572/35x and ET632/60m. β cells were identified by the expression of mCherry. Baseline-normalized cytosolic calcium was quantified using Nikon Elements and GraphPad Prism software.

## Supplemental Figures

**Supplemental Video 1: Control islets show highly synchronized Ca^2+^ oscillations**. Intravital time course video of an islet within the *in vivo* pancreas of a control β cell lineage traced mouse infected with *AAV8-Ins1-GCaMP6s*. Lineage traced β cells are marked by mCherry in red and GCaMP6s is shown in green. Mouse was injected IP with glucose, and video was recorded once blood glucose levels reached ^~^300mg/dL. Z-stack of 8 slices each 8μm apart were recorded at 0.1Hz over 10 minutes. Scale bar is 100μm. Time stamp shown in in upper left corner shows time of image in min:sec.

**Supplemental Video 2: Control islets show highly synchronized Ca^2+^ oscillations**. Intravital time course video of an islet within the *in vivo* pancreas of a control β cell lineage traced mouse infected with *AAV8-Ins1-GCaMP6s*. Lineage traced β cells are marked by mCherry in red and GCaMP6s is shown in green. Mouse was injected IP with glucose, and video was recorded once blood glucose levels reached ^~^300mg/dL. Z-stack of 3 slices each 8μm apart were recorded at 0.2Hz over 12 minutes. Scale bar is 100μm. Time stamp shown in in upper left corner shows time of image in min:sec.

**Supplemental Video 3: *Robo βKO* islets show unsynchronized Ca^2+^ oscillations.** Intravital time course video of an islet within the *in vivo* pancreas of a *Robo βKO* β cell lineage traced mouse infected with *AAV8-Ins1-GCaMP6s*, and retro-orbitally injected with rhodamine-dextran to mark vasculature. Lineage traced β cells are marked by mCherry in red and GCaMP6s is shown in green, and vasculature is shown in yellow. Mouse was injected IP with glucose, and video was recorded once blood glucose levels reached ^~^300mg/dL. Z-stack of 12 slices each 8μm apart were recorded at 0.03Hz over 10 minutes. Scale bar is 100μm. Time stamp shown in in upper left corner shows time of image in min:sec.

**Supplemental Video 4: *Robo βKO* islets show unsynchronized Ca^2+^ oscillations.** Intravital time course video of an islet within the *in vivo* pancreas of a *Robo βKO* β cell lineage traced mouse infected with *AAV8-Ins1-GCaMP6s*. Lineage traced β cells are marked by mCherry in red and GCaMP6s is shown in green. Mouse was injected IP with glucose, and video was recorded once blood glucose levels reached ^~^300mg/dL. Z-stack of 8 slices each 8μm apart were recorded at 0.1Hz over 10 minutes. Scale bar is 100μm. Time stamp shown in upper left corner shows time of image in min:sec.

**Supplemental Video 5: *Robo βKO* islets show unsynchronized Ca^2+^ oscillations.** Intravital time course video of an islet within the *in vivo* pancreas of a *Robo βKO* β cell lineage traced mouse infected with *AAV8-Ins1-GCaMP6s*. Lineage traced β cells are marked by mCherry in red and GCaMP6s is shown in green. Mouse was injected IP with glucose, and video was recorded once blood glucose levels reached ^~^300mg/dL. Z-stack of 3 slices each 8μm apart were recorded at 0.2Hz over 10 minutes. Scale bar is 100μm. Time stamp shown in upper left corner shows time of image in min:sec.

**Supplemental Video 6: A subset of *Robo βKO* islets retain synchronized Ca^2+^ oscillations**. Intravital time course video of an islet within the *in vivo* pancreas of a *Robo βKO* β cell lineage traced mouse infected with AAV8-Ins1-GCaMP6s. Lineage traced β cells are marked by mCherry in red and GCaMP6s is shown in green. Mouse was injected IP with glucose, and video was recorded once blood glucose levels reached ^~^300mg/dL. Z-stack of 8 slices each 8μm apart were recorded at 0.1Hz over 10 minutes. Scale bar is 100μm. Time stamp shown in in upper left corner shows time of image in min:sec.

**Supplemental Figure 1:**
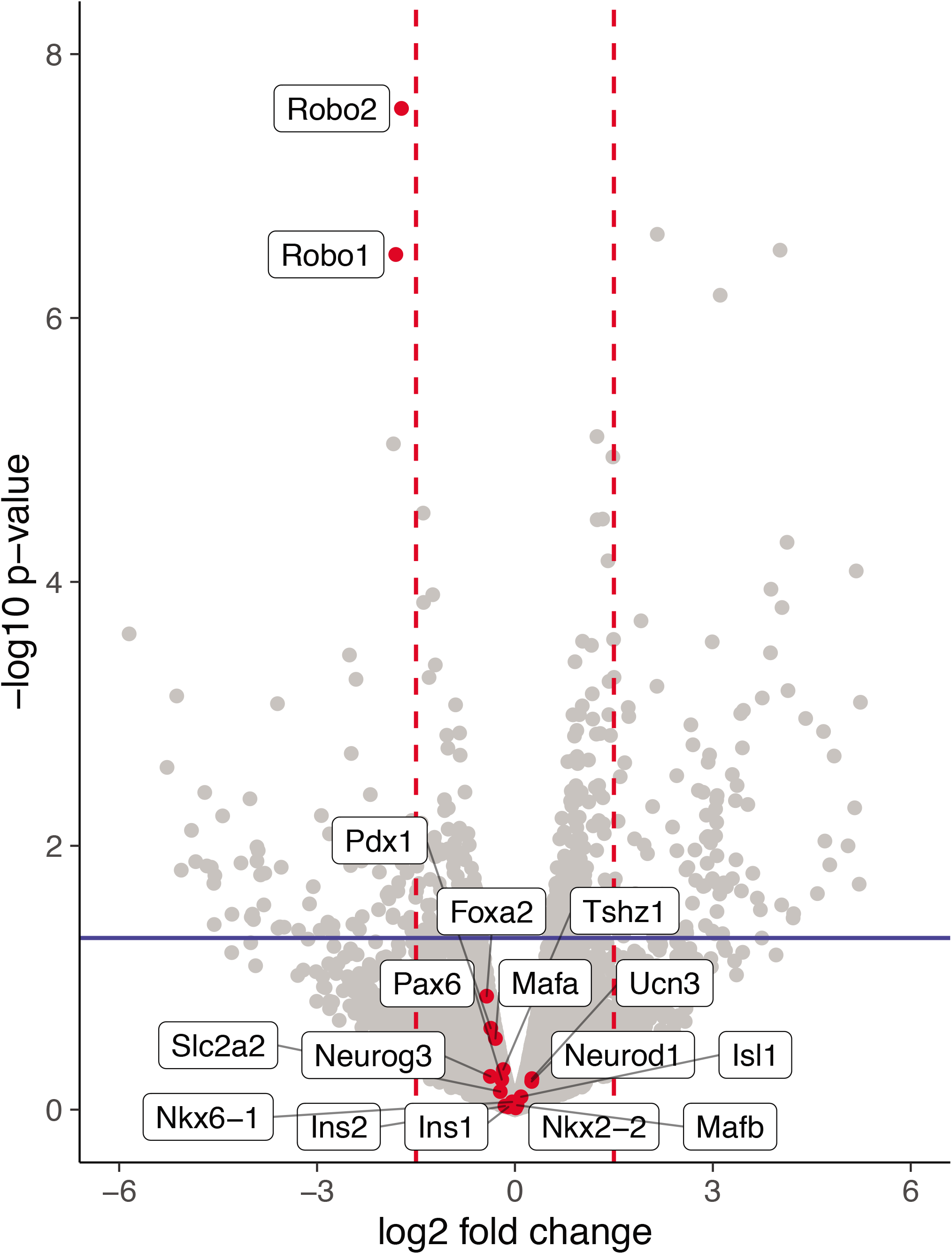
*Robo βKO* islets retain β cell differentiation and maturity markers. Volcano plot of differential gene expression from bulk RNA sequencing on lineage traced FACS sorted β cells from *Robo βKO* and control mice showing no significant differential gene expression of markers (*n*=2 mice from each group) Red lines denote a fold change of 1.5 and blue line denotes a p value of 0.05.

**Supplemental Figure 2:**
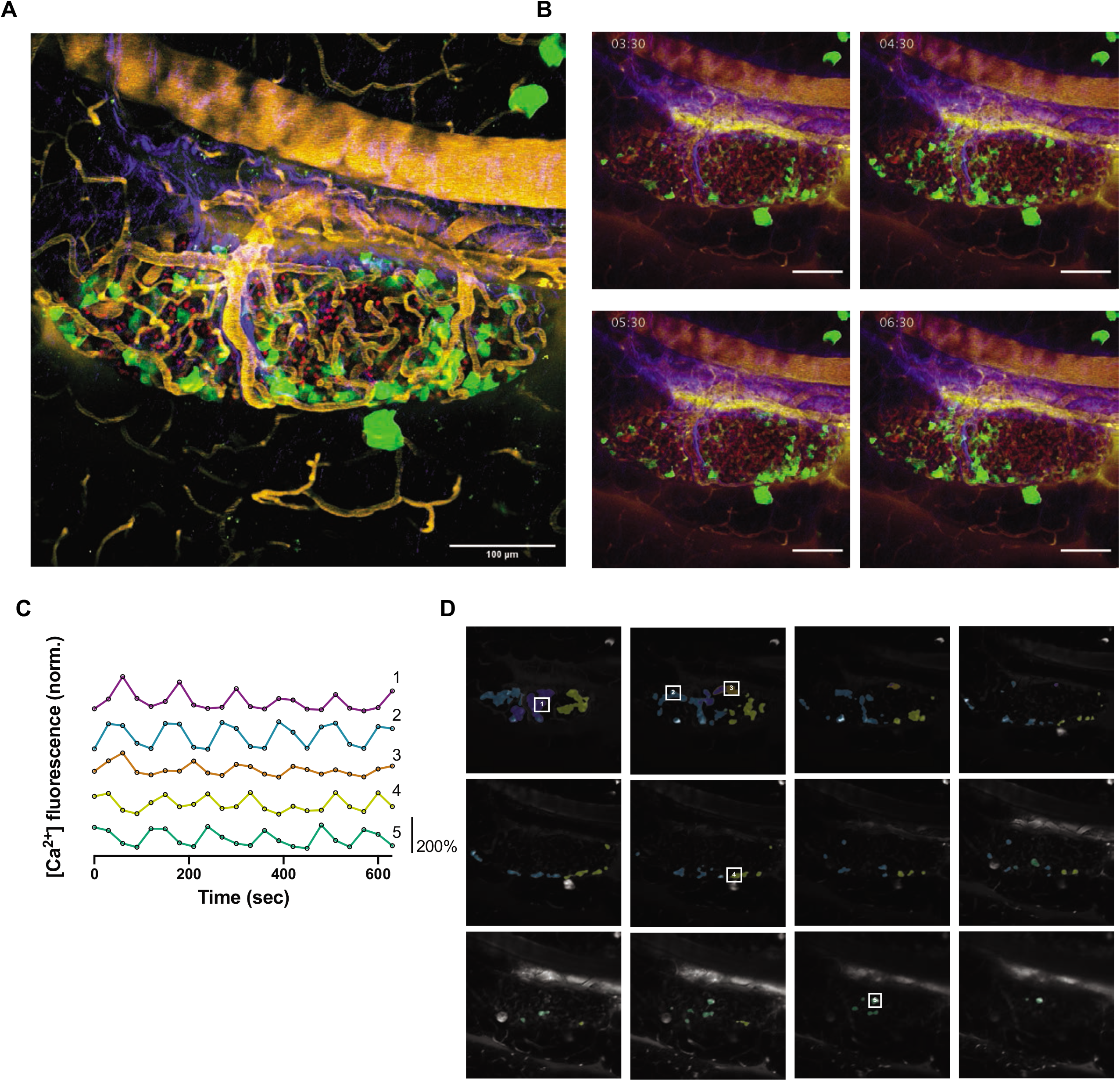
*Robo βKO* islets show uncoordinated whole islet Ca^2+^ oscillations. (A) High resolution maximum intensity projection of a *Robo βKO* islet *in vivo* in an *AAV8-RIP-GCaMP6s*-injected mouse showing GCaMP6s in green, nuclear mCherry β cell lineage tracing in red, and collagen in blue. (B) Stills over one oscillation period from *Robo βKO* islet in supplementary video 3, starting after blood glucose level reached ^~^300mg/dL from IP glucose injection. Video was recorded for 10 minutes with an acquisition speed of 0.03Hz. (C) Representative time courses of Ca^2+^ activity in 4 individual areas from *Robo βKO* islet in supplementary video 3, showing correlation of 43.6% of the active islet area. Time courses are normalized to average fluorescence of individual area over time. Similar color indicates that the time courses have a Pearson’s correlation coefficient of ≥0.75 and matches the region of coordination that is seen in D. (D) False color map of top five largest coordinated areas across z-stack of *Robo βKO* islet from analysis in C. Areas in grey are not coordinated. The color represents a region of coordination with Pearson’s Correlation Coefficient ≥0.75 of GCaMP6s activity. Cells used in time courses in C are labeled.

**Supplemental Figure 3:**
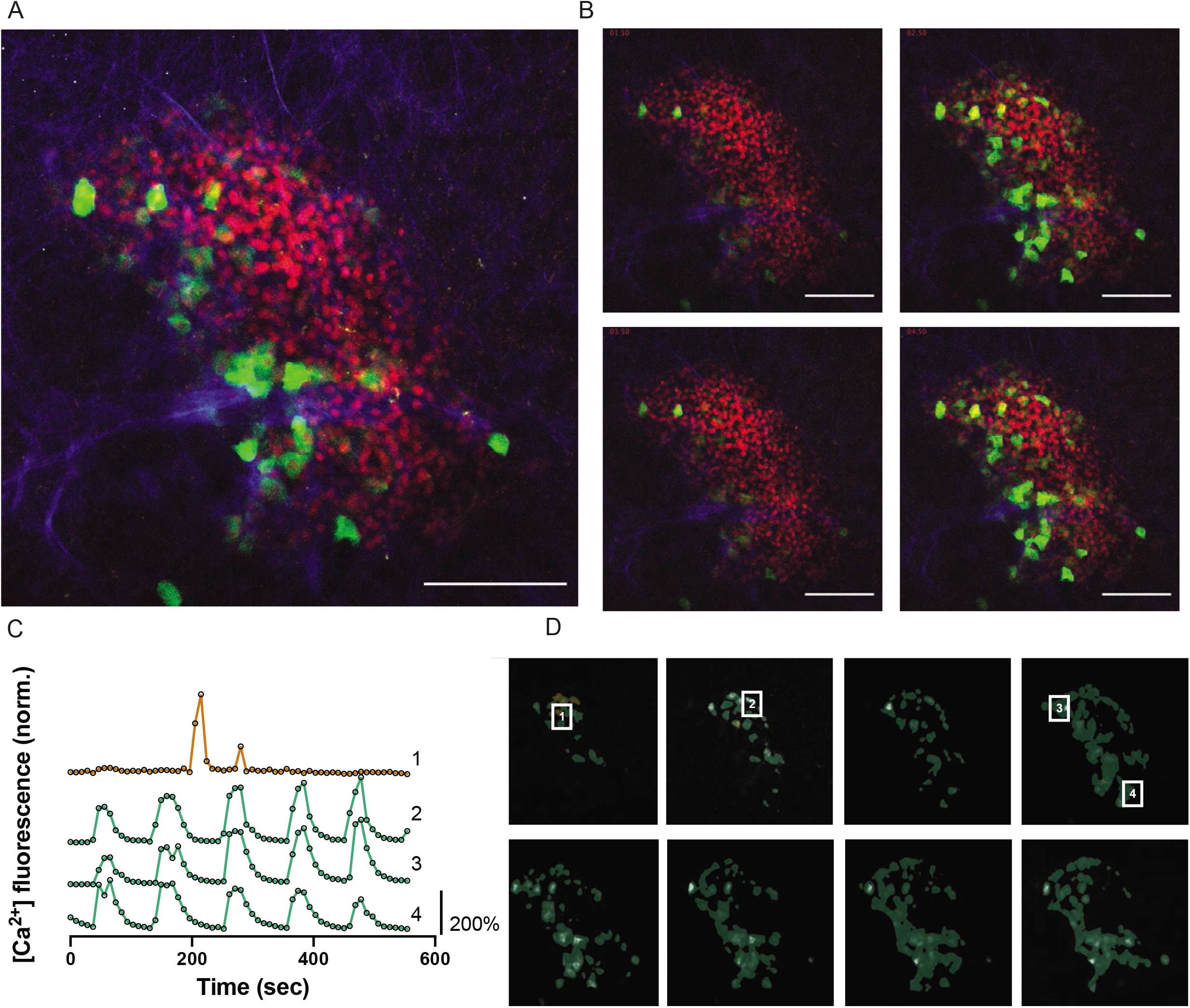
A subset of *Robo βKO* islets show coordinated whole islet Ca^2+^ oscillations. (A) High resolution maximum intensity projection of a *Robo βKO* islet *in vivo* in an *AAV8-RIP-GCaMP6s-injected* mouse showing GCaMP6s in green, nuclear mCherry β cell lineage tracing in red, and collagen in blue. (B) Stills over one oscillation period from *Robo βKO* islet in supplementary video 6, starting after blood glucose level reached ^~^300mg/dL from IP glucose injection. Video was recorded for 10 minutes with an acquisition speed of 0.1Hz. (C) Representative time courses of Ca^2+^ activity in 4 individual areas from *Robo βKO* islet in supplementary video 6, showing correlation of 98% of the active islet area. Time courses are normalized to average fluorescence of individual area over time. Similar color indicates that the time courses have a Pearson’s correlation coefficient of ≥0.75 and matches the region of coordination that is seen in D. (D) False color map of top five largest coordinated areas across z-stack of *Robo βKO* islet from analysis in C. Areas in grey are not coordinated. The color represents a region of coordination with Pearson’s correlation coefficient of ≥0.75 of GCaMP6s activity.

## Notes

### Competing Interest Statement

The authors have declared no competing interest.

